# CD47 Blockade Reprograms the Monocyte-Macrophage Axis to Promote Inflammation Resolution in Atherosclerosis

**DOI:** 10.64898/2026.04.24.720546

**Authors:** Murat Kirtay, Mikael Ispirjan, Benjamin Bonnard, Anna-Lena Bruggner, Lena Boehringer, Moritz Miessler, Norbert Frey, Nicholas J. Leeper, Kai-Uwe Jarr

## Abstract

Atherosclerosis is a chronic inflammatory disease and a leading cause of cardiovascular mortality worldwide. Disease progression is closely linked to defective efferocytosis, the impaired clearance of apoptotic cells, which drives necrotic core expansion and perpetuates arterial inflammation. Targeting the CD47-SIRPα innate immune checkpoint, a dominant “don’t-eat-me” signal, limits atherosclerosis in preclinical models and retrospective human studies. However, how pro-efferocytic intervention reshapes the immune landscape of established atherosclerotic lesions remains incompletely understood.

Here, using single-cell transcriptomics, monocyte fate mapping, and functional analyses across complementary preventive and interventive murine atherosclerosis models, we demonstrate that CD47 blockade fundamentally reprograms the myeloid landscape of established lesions. Anti-CD47 therapy selectively suppresses inflammatory Ly6C^hi^ monocyte recruitment and reduces local macrophage proliferation without altering overall plaque macrophage burden, indicating a qualitative rather than quantitative remodeling of the infiltrate. Concurrently, therapy enriches macrophage subsets bearing pro-efferocytic and macrophage survival-associated transcriptional programs, restoring defective apoptotic cell clearance in situ. Cross-species integration with an independent human coronary artery single-cell dataset identifies a conserved TREM2^hi^ macrophage population that natively harbors the efferocytosis machinery reactivated by therapy in mice.

Together, these findings demonstrate that innate immune checkpoint inhibition by CD47 blockade drives a coordinated reprogramming of monocyte–macrophage dynamics, simultaneously suppressing inflammatory influx and enriching efferocytic capacity. This dual mechanism advances our understanding of how pro-efferocytic therapies resolve vascular inflammation in atherosclerosis.

## Introduction

Atherosclerosis is a lipoprotein-driven chronic inflammatory disease. While lipid-lowering therapies, such as statins and PCSK9 inhibitors^1^, have improved clinical outcomes, residual cardiovascular risk remains, largely driven by unresolved arterial inflammation^2,3^. Although atherosclerotic plaques harbor a highly complex immune microenvironment involving both adaptive (e.g., T and B cells) and innate immune populations^4–6^, plaque-associated macrophages occupy a central position in driving disease progression^7^. Originating primarily from functionally distinct circulating monocyte subsets (such as pro-inflammatory Ly6C^hi^ and patrolling Ly6C^lo^ monocytes)^8–10^, these cells govern plaque composition. Beyond aberrant lipid uptake and processing, plaque macrophages exhibit impaired clearance of apoptotic cells, a process termed efferocytosis^11–15^. Defective efferocytosis drives necrotic core expansion, destabilizes plaques, and perpetuates local inflammation^11,16,17^. Therefore, restoring innate immune homeostasis and macrophage function represents a prime therapeutic target.

A key driver of this impaired clearance is the upregulation of Cluster of Differentiation 47 (CD47), a “don’t-eat-me” signal that interacts with Signal-Regulatory Protein alpha (SIRPα) on myeloid cells to inhibit phagocytosis^18–20^. Targeting this innate immune checkpoint has emerged as a promising strategy: anti-CD47 antibodies reactivate efferocytosis, reduce lesion burden, and have been associated with reduced vascular inflammation in both preclinical and retrospective human studies^20–25^. Notably, while landmark clinical trials like CANTOS (targeting IL-1β via canakinumab) validated the inflammatory hypothesis of atherosclerosis, they also highlighted the infectious risks of broad immunosuppression^26^. This underscores the urgent need for therapies that do not merely suppress inflammation but actively induce resolution and tissue repair.

While the CD47-SIRPα axis represents the only pro-efferocytic strategy to have transitioned to clinical testing in atherosclerotic cardiovascular disease^27^, a critical knowledge gap remains. Despite evidence that CD47 blockade enhances efferocytosis, it is unknown whether this therapy solely augments immediate phagocytic activity or whether it fundamentally reprograms the underlying inflammatory monocyte–macrophage axis. The precise mechanisms by which this checkpoint blockade alters the local immune landscape remain undefined.

Here, we combine single-cell transcriptomics with functional analyses in murine atherosclerosis and cross-species validation in human coronary plaques to define how pro-efferocytic therapy remodels the innate immune compartment. We demonstrate that CD47 blockade reshapes the immune landscape toward a pro-resolution state by modulating the monocyte-macrophage axis, reactivating a conserved efferocytosis-competent macrophage program, and altering inflammatory recruitment and local proliferation. Furthermore, we identify a homologous macrophage population in human plaques that shares this machinery, supporting the translational relevance of macrophage reprogramming as a therapeutic mechanism of anti-CD47 therapy. Together, these findings advance our mechanistic understanding of pro-efferocytic therapy and provide a translational framework to inform the clinical development of pro-efferocytic treatments.

## Methods

### Data Availability

Extensive methods, including statistical analysis and the Major Resources Table, are included in the Supplemental Material. All raw data and analytical methods are available from the corresponding author upon reasonable request. Sequencing data have been deposited at the public Gene Expression Omnibus under accession number GSE319762.

### Mouse Studies

All mice were housed in a pathogen-free animal facility and maintained on a 12-h light:dark cycle at 22 °C with free access to food and water. Mice were acclimatized for 1–2 weeks before experiments. Experiments were performed with 6- to 8-week-old male and female *Ldlr^-/-^* mice (B6.129S7-*Ldl*r^tm1Her^/J, strain #:002207), unless indicated otherwise. For the preventive and interventive therapy model, both male and female mice were included in functional, histological, and flow cytometric analyses. Single-cell RNA-sequencing was performed exclusively in male mice to minimize transcriptional variance attributable to sex as a confounding variable in this single-library-per-group design. A total of 90 mice were used for the present study. The animals were assigned to experimental groups by complete randomization and all analyses were blinded whenever possible through numerical coding of sample identification. For earlier power analysis, the following parameters were used based on previous experience with the therapeutic agents and similar atherosclerotic models^20,21,28,29^: α of 0.05; power of 0.80. For the preventive therapy model, mice were fed a high-fat, high-sucrose Western Diet^30–33^ (WD, 21 % butter fat and approx. 0.21 % cholesterol, 34 % sucrose, corresponding to TD.88137, ssniff) and injected simultaneously with either the inhibitory anti-CD47 (BioXCell, clone MIAP410^20,29^) or the respective IgG isotype control (BioXCell, clone MOPC-21) intraperitoneally every other day (200 µg per dose) for 12 weeks. For the interventive therapy model, the mice were fed the same WD for 14 weeks and the therapy was applied for the last 6 weeks of WD feeding. All animal studies were approved by the local Animal Care and Use Committees (Regierungspräsidium Karlsruhe, AZ: G-125/22) and conformed to the guidelines from Directive 2010/63/EU of the European Parliament on the protection of animals used for scientific purposes.

### Aortic Single-cell Preparation and CD45^+^ Leukocyte Isolation

Male mice (n=4 IgG; n=4 anti-CD47), after 12 weeks of WD feeding (approximately 18-20 weeks of age), were used to isolate viable CD45^+^ aortic leukocytes. Male mice were selected for scRNA-seq to reduce sex-related transcriptional heterogeneity within the pooled single-library design; the functional and histological experiments include both sexes as described. Five minutes before euthanasia (via anesthesia overdose) and aorta isolation, the mice were injected i.v. with 2 µg CD45-BV510 antibody to stain circulating leukocytes. Intracardiac perfusion was performed with ice-cold PBS to flush out the blood. Aortas were isolated by cleaning perivascular adipose tissue and stored in PBS until digestion. Aortas from the abovementioned mice were individually digested for 40 min at 37 °C in RPMI medium containing 450 U/ml Collagenase I (Sigma-Aldrich, C0130), 125 U/ml Collagenase XI (Sigma-Aldrich, C7657), and 60 U/ml Hyaluronidase (Sigma-Aldrich, H3506), as described previously^4^. The digested tissue was meshed through a 70-µm cell strainer to obtain a single-cell suspension. The cells were centrifuged at 500 x g for 5 min at 4°C and were resuspended in FACS Buffer (2 % FCS, 2 mM EDTA in PBS) containing TruStain FcX anti-mouse CD16/CD32 (Biolegend, 101320, 1:100) to block non-specific binding of immunoglobulin to Fc receptors and incubated for 10 min at 4 °C. The cells were subsequently stained with SYTOX Green viability dye (L/D), TER119-BV510, and CD45-PerCP Cy5.5 for 30 min at 4 °C. Due to the low yield of viable CD45^+^ leukocytes recoverable from individual aortas, aortas from each group were pooled (n = 4 per group, yielding n = 1 sequencing library per group) before sorting of viable SYTOX Green^-^TER119^-^ CD45^+^ cells using a 100-µm nozzle by excluding CD45-BV510^+^ circulating leukocytes. This pooling strategy, which is an established approach in the field for low-input aortic immune profiling^4,34,35^, precludes formal statistical replication at the single-cell sequencing level and was therefore complemented by independent biological replicates in all functional, flow cytometric, and histological analyses. Sorted cells were immediately processed for single-cell RNA-sequencing.

### Single-cell RNA-sequencing

Samples were resuspended to a concentration of 600–1,000 cells/μl and loaded into the 10x Chromium system to generate single-cell barcoded droplets using the Single Cell 3’ Solution v3 Reagent Kit (10x Genomics) following the manufacturer’s specifications. Generated libraries were sequenced using a NextSeq 2000 Sequencing System in paired-end mode to reach approximately 200,000 reads per single cell.

### Mouse Single-cell RNA-sequencing Data Analysis

Mouse single-cell RNA-sequencing data were preprocessed using 10x Cell Ranger cloud software (Cell Ranger v9.0.1), which included data demultiplexing, barcode processing, alignment, and single-cell 3′ gene counting. The reads mapped to reference mouse genome (UCSC mm10) were used to generate a gene-barcode matrix for downstream analysis. Cells with < 200 detected genes, < 500 read counts, or a percentage of mitochondrial genes > 5 % were filtered out. Cells not explicitly expressing *Ptprc* (encoding for CD45) were excluded from the analysis. The resulting data were log-normalized, scaled, and regressed on the number of unique molecular identifiers per cell and the percentage of mitochondrial gene content. The datasets from anti-CD47 and IgG-treated animals, together with the *Ldlr^-/-^* datasets from Cochain et al. (GSE97310), were integrated for batch correction using Harmony^36^. Data visualization, detection of marker genes, gene expression analyses, trajectory mapping of monocytes/macrophages using Monocle 3, and CellChat analyses were conducted using BioTuring BBrowser Pro^37^. Differentially expressed genes with an FDR < 0.1 based on the Wilcoxon rank-sum test with Benjamin-Hochberg correction were considered statistically significant.

### Human Single-cell RNA-sequencing Data Analysis

Human single-cell RNA-sequencing data of human atherosclerotic coronary arteries were obtained from the Gene Expression Omnibus (GSE131778)^38^ and analyzed using BioTuring BBrowser Pro^37^. Cells with < 200 detected genes, < 500 read counts, or a percentage of mitochondrial genes > 5 % were filtered out. A total of 11,756 cells were included in the analysis. Data were log-normalized before dimensional reduction. Cell clustering of the entire cell set using the Louvain method was performed at a 0.3 resolution. To analyze the macrophages, myeloid cells were annotated based on marker gene expression, subsetted, and reclustered using a resolution of 0.5^38^.

### Tissue Preparation, Blood Analysis, Immunofluorescence, and Histological Analyses

Tissue preparation and histological analyses were performed as previously described^21,28^. After euthanasia, blood samples were rapidly collected. Mice were then perfused with ice-cold PBS via cardiac puncture, and spleen, aorta-draining lymph nodes, bone marrow, and aorta were isolated. Serum was obtained by centrifuging the blood in clotting activator-containing micro-sample tubes at 2,000 x g for 10 min at 4°C. To confirm the induction of the pro-atherogenic metabolic phenotype, circulating lipids (total cholesterol, LDL, and HDL) and non-fasting glucose were assessed by the Heidelberg University Hospital Diagnostic Laboratory using an automated clinical chemistry analyzer (Roche Diagnostics). For analysis of aortic roots, hearts were fixed in 4 % paraformaldehyde (PFA) and subsequently cryoprotected in 30 % sucrose solution before being embedded in optimal cutting temperature (OCT) compound. Aortic root sections at 8 µm thickness were prepared using a cryostat (Leica CM 1950). To accurately capture the entire plaque volume, four step-sections per mouse were collected at 80-µm intervals, beginning from the anatomical base of the aortic root (identified by the appearance of the aortic valve leaflets). Plaque lipid area (in mm^2^) was quantified by Oil Red O staining (Sigma-Aldrich, O1516). Because the external elastic lamina (EEL) is not readily visible with ORO alone, sections were counterstained with Hematoxylin. Total vessel area was calculated by encircling the EEL on these sections, which then served as the anatomical reference to determine the percentage of plaque burden. For immunofluorescence staining of atherosclerotic lesions, cryosections were blocked using 5 % BSA in PBS. Next, sections were incubated overnight at 4 °C with the following primary antibodies: CD68 (Invitrogen, 14-0681-82, 1:200), cleaved caspase-3 (Cell Signaling Technology, 9661, 1:200), and Ki67 (Abcam, ab16667, 1:50). After extensive washing, sections were incubated with secondary antibodies from Thermo Fisher Scientific: Alexa Fluor-647 goat anti-rat (A32733, 1:400) and Alexa Fluor-555 goat anti-rabbit (A48263, 1:400).

Counterstaining to visualize nuclei was performed by incubating with DAPI. Both histological sections and immunofluorescence staining sections were imaged with a 20x objective using a Zeiss Axioscan 7. Sections were analyzed using QuPath software (v.0.6.0) in a blinded fashion.

### Flow Cytometry

Blood was filtered through a 30-µm cell strainer, washed with FACS Buffer (2 % FCS, 2 mM EDTA in PBS), and erythrocytes were lysed two times with ACK Lysing Buffer (Gibco, A1049201) for 5 min at RT. Spleens were mechanically disrupted on a 40-µm cell strainer and subjected to erythrocyte lysis before staining. Femurs and tibias were cut open, and bone marrow was isolated by centrifuging the bones at 15,000 x g for 30 sec. For flow cytometry staining, all prepared cells from different tissues were incubated in TruStain FcX anti-mouse CD16/CD32 (Biolegend, 101320, 1:100) for 10 min at 4 °C to block non-specific binding of immunoglobulin to Fc receptors. After blocking, cells were stained with a fluorophore-conjugated antibody cocktail (Table S1) containing a fixable live/dead dye (eBioscience™ Fixable Viability Dye eFluor™ 660, Thermo Scientific) for 30 min at 4 °C, protected from light. After being washed with FACS Buffer, samples were either analyzed directly or fixed in IC Fixation Buffer (eBioscience™) for 20 min at 4 °C, protected from light to preserve fluorescence signal prior to acquisition. Data were acquired with a BD FACSCelesta™ Cell Analyzer and processed in FlowJo v10 (TreeStar).

### Differential Labelling and Tracking of Blood Monocytes

To assess monocyte recruitment specifically within the interventive therapy model, circulating Ly6C^lo^ monocytes were labeled by i.v. injection of 1 µm Fluoresbrite YG microspheres (Polysciences, Inc, PA) diluted 1:4 in sterile PBS, 3 days before sacrifice, as described^35,39^. To track the recruitment of newly generated Ly6C^hi^ monocytes to the plaques, their proliferating bone marrow progenitors were pulse-labeled by an i.p. injection of 1 mg of EdU (5-ethynyl-2’-deoxyuridine, Thermo Scientific) 3 days before sacrifice to analyze recruitment to the plaques, as described^35,39^. The efficiency of labeling was verified 24 h after injection by flow cytometry analysis of blood monocytes. EdU labeling was determined using the Click-IT EdU Pacific Blue Flow Cytometry Assay Kit (Invitrogen, C10418). The EdU-labeled monocytes in aortic root sections were analyzed using the Click-iT™ Plus EdU Cell Proliferation Kit for Imaging, Alexa Fluor™ 647 dye (Invitrogen, C10640). At least 3-4 sections per animal were analyzed and average EdU^+^ and bead^+^ cell quantification per section were reported in the figures.

### RNA isolation and quantitative Polymerase Chain Reaction (qPCR)

Aortic arches containing atherosclerotic plaques were homogenized in TRIzol Reagent (Thermo Fisher Scientific), and total RNA was isolated from homogenized aortas. Briefly, 200 µl chloroform was added per 1 ml TRIzol homogenate, and the upper aqueous phase containing RNA was isolated after centrifuging the sample at 12,000 x g for 15 min at 4 °C. RNA was precipitated in an equal volume of isopropanol containing GlycoBlue (Invitrogen, AM9515). After ethanol washes, the RNA pellet was resuspended in DNase/RNase-free water. RNA concentration and purity were quantified with NanoDrop One (Thermo Fisher Scientific). An equal amount of total RNA from each sample was reverse transcribed using iScript™ cDNA Synthesis Kit (BioRad, 1708891), and qPCR of the cDNA samples was performed using iQ™ SYBR® Green Supermix (BioRad, 1708882) on a QuantStudio 5 (Applied Biosystems). Data were quantified by the 2^−ΔΔCt^ method and normalized to *Gapdh* as an internal control. The stability of *Gapdh* expression across all experimental conditions was explicitly confirmed to ensure reliable normalization. The qPCR primers used in the study are listed in Table S2.

### Statistical Analysis

Statistical analyses were performed using GraphPad Prism 10 (GraphPad Inc.). Data and error bars present mean ± SD for parametric and median ± IQR for non-parametric results. Normality of data was determined by performing a Shapiro–Wilk normality test (α = 0.05). Normally distributed data were analyzed using an unpaired Student’s *t*-test (two-tailed) or one-way analysis of variance (ANOVA) with Tukey’s multiple comparisons test or two-way ANOVA with Geisser-Greenhouse correction. For samples that had unequal variances (determined by F-test), an unpaired Welch’s t-test (two-tailed) was used. If the data set did not pass the normality test (p-value < 0.05), the statistical significance was determined by Mann Whitney U test (two-tailed) or Kruskal-Wallis test, followed by Dunn’s multiple comparisons test for more than two groups. Exact p-values are indicated. P values for the differentially expressed genes were adjusted for multiple testing using the Benjamin-Hochberg FDR. All data behind the statistical analysis and all p values are provided in Source Data.

**Table S1.** FACS Antibodies.

| **Antigen** | **Fluorophore** | **Clone** | **Provider** | **Catalog Nr.** |
| --- | --- | --- | --- | --- |
| CD45 | PerCP Cy5.5 | 30-F11 | BD Biosciences | 561869 |
| TER-119 | BV510 | TER-119 | Biolegend | 116237 |
| CD45 | BV510 | 30-F11 | Biolegend | 103138 |
| CD11b | Pacific Blue | M1/70 | Biolegend | 101224 |
| Ly6C | PE | HK1.4 | Biolegend | 128008 |
| Ly6G | PE-Cy5 | 1-A8 | Biolegend | 127672 |
| F4/80 | PE-Cy7 | BM8 | Biolegend | 123114 |
| CD11c | BV605 | N418 | Biolegend | 117333 |
| CX3CR1 | BV711 | SA011F11 | Biolegend | 149031 |
| CD45 | APC-Cy7 | 30-F11 | Biolegend | 103116 |
| CD11c | BV510 | N418 | Biolegend | 117353 |
| Ly6G | BV605 | 1-A8 | Biolegend | 127639 |
| CD115 | PE-Cy7 | AFS98 | Biolegend | 135523 |

**Table S2.**
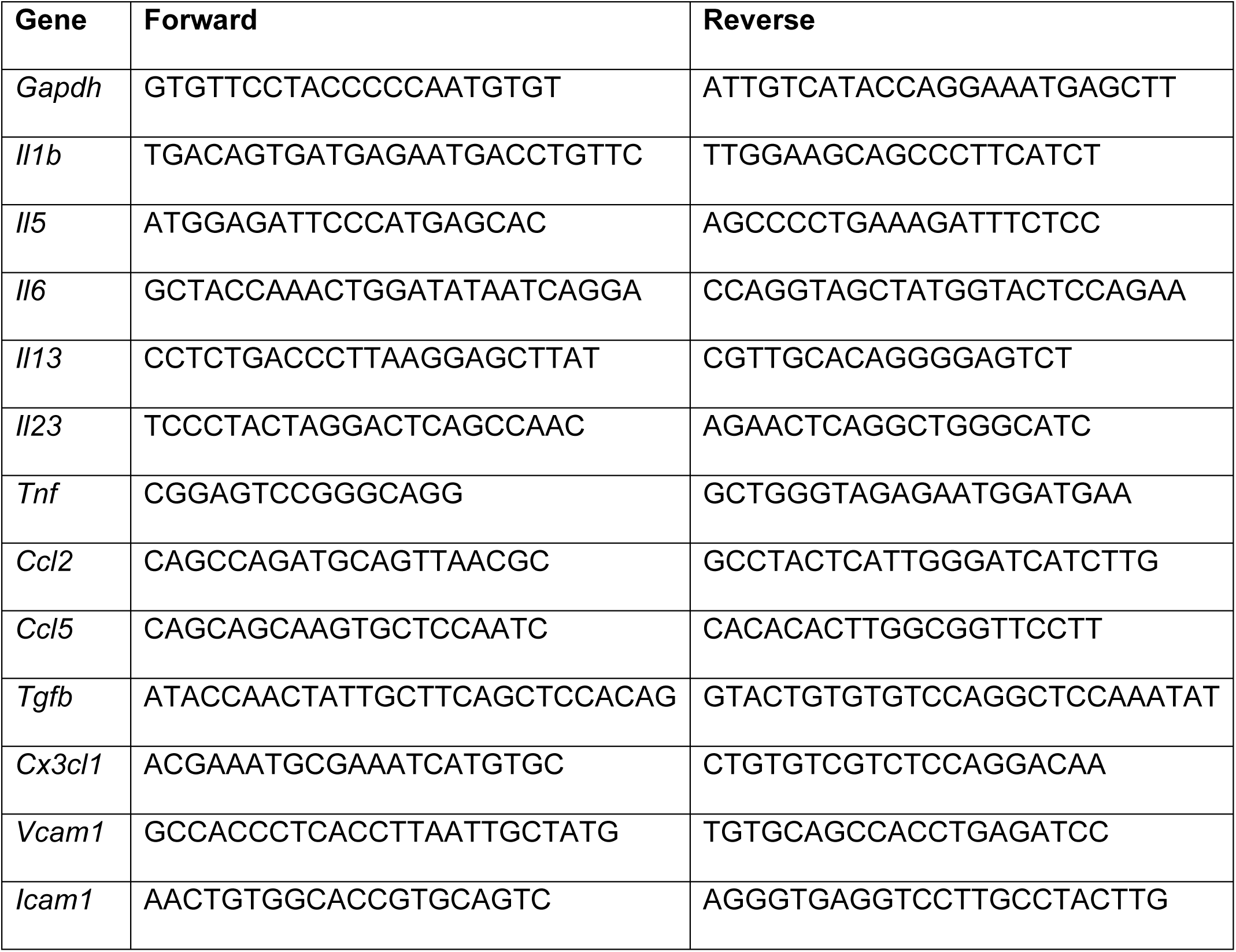
Primer sequences.

## Results

### Anti-CD47 Treatment Reprograms the Aortic Myeloid Landscape

Blockade of the CD47-SIRPα axis promotes atherosclerotic plaque regression and resolves vascular inflammation across murine models, large animals, and human cohorts^20–23,29^. However, the cellular mechanisms driving these therapeutic effects remain incompletely understood. To map the immunomodulatory landscape of CD47 blockade in a highly translationally relevant setting, *Ldlr^-/-^* mice were fed a high-fat, high-sucrose Western Diet for 12 weeks^30,31^. We then performed single-cell RNA-sequencing (scRNA-seq) on viable CD45^+^ aortic leukocytes from *Ldlr*^-/-^ mice in a prevention setting, treating the mice concurrently with an anti-CD47 antibody or an IgG isotype control for 12 weeks (Figure 1A and Figure S1A). Of note, anti-CD47 treated animals did not show any differences in their body and heart weight but had an increased spleen weight as reported previously (Figure S1B–S1D)^20,24^. To confirm the induction of the pro-atherogenic metabolic phenotype, circulating lipids and non-fasting glucose were compared between the cohorts (Figure S1E). We integrated 2,854 high-quality cells from our experimental cohorts (1,237 IgG; 1,617 anti-CD47) with 1,315 reference cells from a publicly available scRNA-seq dataset of *Ldlr*^-/-^ mice under chow diet (CD, n = 381) and Western Diet (WD, n = 934)^4^, yielding a consolidated atlas of 4,169 cells after batch correction (Figure 1B; Figure S1F and S1G). Unsupervised clustering based on canonical marker genes (e.g., *Apoe, Cd3e, Cd209a, Trem2)* revealed seven major immune cell lineages (Figure 1B and 1C). While the IgG control group mirrored the atherogenic WD phenotype, anti-CD47 treatment shifted the mononuclear phagocyte (MPC) and antigen-presenting cell (APC) compositions toward a non-atherogenic baseline (Figure 1D). To resolve myeloid heterogeneity, we reclustered the MPC and APC compartments (macrophages, monocytes and monocyte-derived dendritic cells, MoDCs), identifying 11 transcriptionally distinct subsets. These included established atherosclerosis-associated populations^4,5^, such as TREM2^hi^ foamy macrophages (*Trem2, Spp1, Cd9*), inflammatory macrophages (*Cxcl2, Nlrp3, and Nfkbiz*) and tissue-resident macrophages (*Cd163, Gas6, and Mrc1*) (Figure 1E and 1F). Consistent with myeloid reprogramming, anti-CD47 treatment was associated with a reduction in TREM2^hi^ metabolic macrophages in this prevention model, while homeostatic resident-like populations enriched in phagocytosis-related gene expression appeared preserved^4^ (Figure 1G). Collectively, these compositional shifts are consistent with CD47 blockade driving myeloid reprogramming toward a more reparative macrophage phenotype during atherogenesis.

**Figure 1.**
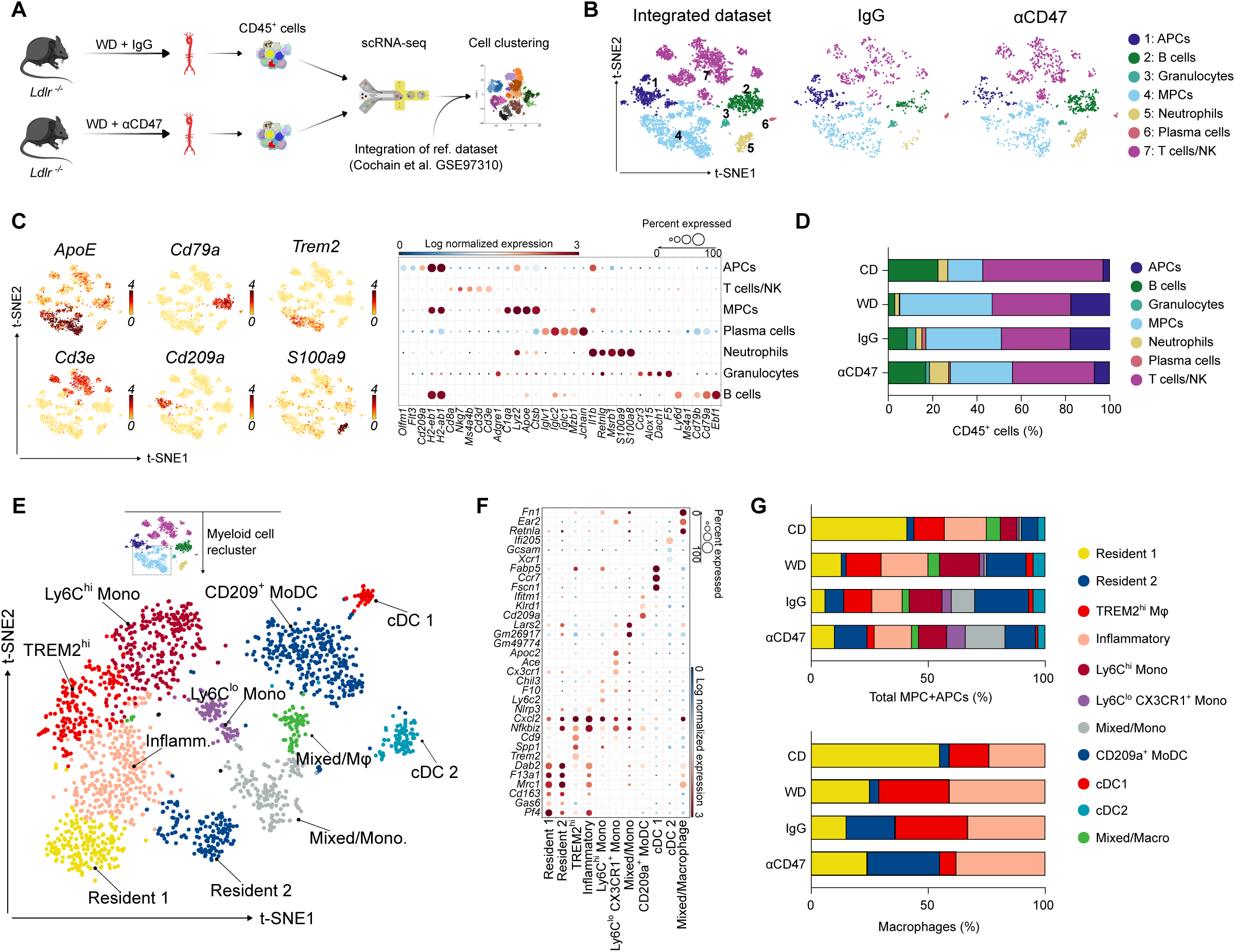
Single-cell RNA-sequencing reveals remodeling of TREM2^hi^ macrophages upon CD47 blockade. A, Schematic of CD45^+^ leukocyte isolation from aortic single-cell suspensions followed by single-cell RNA-sequencing (scRNA-seq). WD: Western Diet. **B,** t-distributed stochastic neighbor embedding (t-SNE) plot of aligned gene expression data from single CD45^+^ cells extracted from the aortas of *Ldlr^−/−^* mice. Mice were fed a Western Diet (WD) for 12 weeks and treated with either control IgG (n=1,237 cells) or anti-CD47 antibody (n=1,617 cells) (n=4 mice pooled per group, yielding one sequencing library per group). Publicly available scRNA-seq data from aortic CD45^+^ leukocytes of mice fed a chow diet (CD) and a WD (Cochain et al., 2018) were integrated to facilitate cell cluster alignment. **C,** Expression of 6 selected marker genes per cluster (left) and dot plot of 5 selected marker genes used for cluster identification (right). The color scale represents log-transformed gene expression. **D,** Quantification of 7 major cell populations within the treatment groups. **E,** t-SNE plot of the myeloid cell compartment after subsetting and reclustering from the total CD45^+^ population. **F,** Dot plot of marker gene expression used for identification of mononuclear phagocyte (MPC) and antigen-presenting cell (APC) subsets. 3 selected cluster-defining genes are shown. **G,** Quantification of identified macrophage and dendritic cell populations within the myeloid compartment (top) and macrophage compartment alone (bottom).

### Anti-CD47 Therapy Alters Monocyte Differentiation Trajectories

Given the macrophage reprogramming observed under CD47 blockade, we investigated their monocyte precursors. Mouse monocytes classically divide into pro-inflammatory Ly6C^hi^ and patrolling Ly6C^lo^ subsets, with the latter serving as precursors for resolving macrophages^9,40^. To map the development trajectories of aorta-infiltrating monocytes into macrophages, we performed pseudotime analysis using Monocle 3. By designating monocytes as the root population, trajectory analysis suggested a relative bias away from a foamy macrophage fate (Figure 2A; Figure S2A). Reclustering of the monocyte compartment identified five transcriptionally distinct clusters (Figure 2B; Figure S2B and S2C). Intriguingly, Ly6C^lo^ monocytes expanded in treated aortas, a finding confirmed by flow cytometry (Figure 2C; Figure S3A). Furthermore, treatment increased the frequency of F4/80^+^ Ly6C^lo^ macrophages, a population derived from Ly6C^hi^ monocytes and linked to inflammation resolution^41^, while preserving the total F4/80^+^ macrophage population (Figure 2C). Systemically, while blood Ly6C^hi^ monocyte levels trended downward, their numbers expanded in splenic and lymph node reservoirs. This peripheral redistribution, rather than systemic depletion, is consistent with a sequestration mechanism that limits monocyte availability for plaque recruitment (Figure 2D; Figure S3B). Importantly, we observed no changes in the bone marrow compartment (Figure S3B). To confirm the shift toward resolution, we analyzed inflammatory signatures within the MPC cluster. Anti-CD47 therapy markedly downregulated classical pro-inflammatory genes at the single-cell level, including *Nlrp3, Tnf*, and *Il1b* (Figure 2E), as well as *Nfkb1* and *Nfkbiz* (Figure S3C). At the tissue level, qPCR of the ascending aorta confirmed downregulation of the pro-inflammatory genes *Il1b, Il6, Ccl2* and *Ccl5*, alongside the adhesion molecule *Vcam1* (Figure 2F; Figure S3D). Notably, *Tnf* expression did not reach significance by qPCR despite its reduction in scRNA-seq data, likely reflecting dilution of the macrophage-specific signal within bulk aortic tissue containing multiple non-myeloid cell types. No changes were detected for *Il5, Il13, Il23, Cx3cl1*, or *Icam1* (Figure S3D).

**Figure 2.**
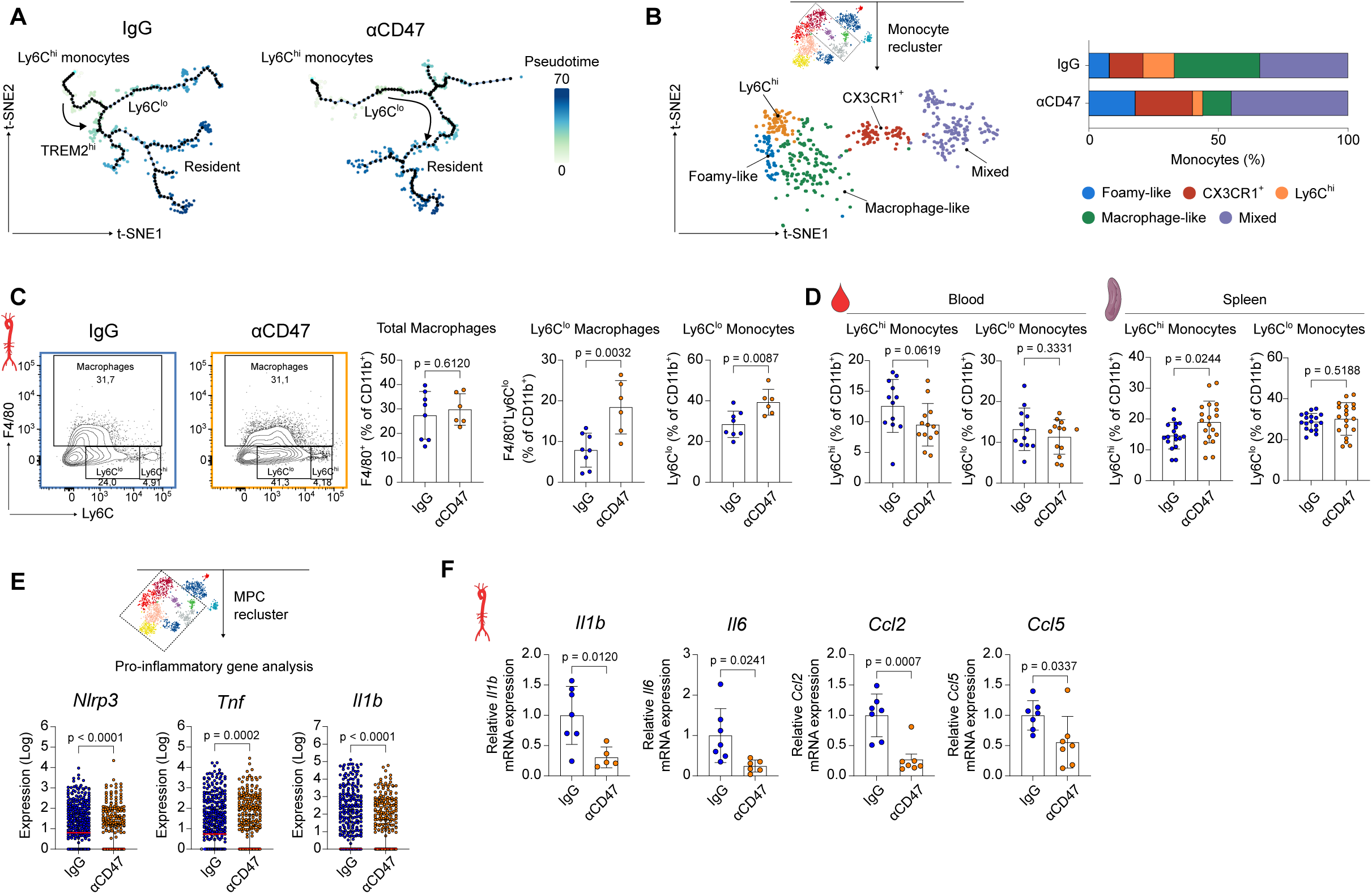
CD47 blockade induces transcriptional and phenotypic rewiring of monocytes. A, Monocle3 pseudotime trajectory of monocyte-macrophage populations from IgG- and anti-CD47-treated animals superimposed on a t-SNE plot. The color scale represents differentiation progression along pseudotime. **B,** t-SNE plot depicting unsupervised clustering of aortic monocytes (left) and proportional representation of each identified subcluster within the total monocyte population (right). **C,** Flow cytometry quantification of F4/80^+^ total macrophages, F4/80^+^ Ly6C^lo^ macrophages, and F4/80^-^ Ly6C^lo^ non-classical monocytes in the aortas of *Ldlr^-/-^* mice after 12 weeks of WD feeding and treatment with IgG (n=8) or anti-CD47 (n=6). **D,** Flow cytometry analysis of Ly6C^hi^ and Ly6C^lo^ monocytes in blood (n=12 IgG; n=13 anti-CD47) and spleen (n=18 IgG; n=19 anti-CD47). **E,** scRNA-seq gene expression of *Nlrp3*, *Tnf,* and *Il1b* in the MPC cluster. **F,** Quantitative polymerase chain reaction (qPCR) analysis of *Il1b*, *Il6*, *Ccl2,* and *Ccl5* expression in aortic arch samples (n=7 IgG; n=5-7 anti-CD47). Data and error bars present mean ± SD. Statistical analysis was performed using unpaired Student’s *t*-test (two-tailed). All data and statistical analysis are provided in Source Data.

### CD47 Blockade in Advanced Atherosclerosis Induces Pro-efferocytic Myeloid Cell Reprogramming

To disentangle direct myeloid reprogramming from secondary changes caused by plaque regression, we treated mice with anti-CD47 antibodies only during the final six weeks of WD-feeding (Intervention model; Figure 3A). Importantly, this regimen did not alter lesion size under the pro-atherogenic metabolic phenotype (Figure 3B; Figure S4A), allowing the assessment of immune dynamics independent of plaque burden. In this established plaque environment, scRNA-seq analysis of the aortic arch revealed a distinct shift in the MPC compartment; while total dendritic cell (DC) populations (expressing *Cd209a, Itgax, Klrd1, Ccr7, Ifi205*) contracted, macrophage subsets expanded (Figure 3C–3F; Figure S4B and S4C). Unlike the prevention model, specific macrophage subsets were not depleted; rather, their transcriptional state was altered. Transcriptional profiling of MPC and APC clusters, alongside global gene expression analysis, revealed that CD47 blockade upregulated a core pro-efferocytic program. This included lipid metabolism genes (*Apoe, Trem2*), complement factors (*C1qa, C1qb, C1qc*), bridging molecules (*Mfge8, Gas6*), the anti-inflammatory cytokine *Il10*, and scavenger receptors *(Mertk, Cd36*), while concomitantly downregulating the “don’t-eat-me” signal *Cd24a*, a phagocytic checkpoint functionally analogous to CD47, which suppresses efferocytosis through interaction with SIGLEC G/SIGLEC-10 on myeloid cells (Figure 4A and 4B; Figure S5A)^42^. Pathway analysis confirmed an enrichment of efferocytosis signaling and macrophage motility (Figure 4C and 4D; Figure S5B–S5D). Validating this functional restoration *in situ*, we observed a 3.5-fold increase in the ratio of CD68^+^ macrophage-associated to free apoptotic cells in treated plaques (Figure 4E). Importantly, the absolute number of apoptotic cells did not differ significantly between groups (Figure S5E), indicating that the improved ratio reflects enhanced macrophage clearance capacity rather than a reduction in apoptotic cell generation per se. These data demonstrate that CD47 blockade reprograms existing macrophages toward a reparative, pro-efferocytic state.

**Figure 3.**
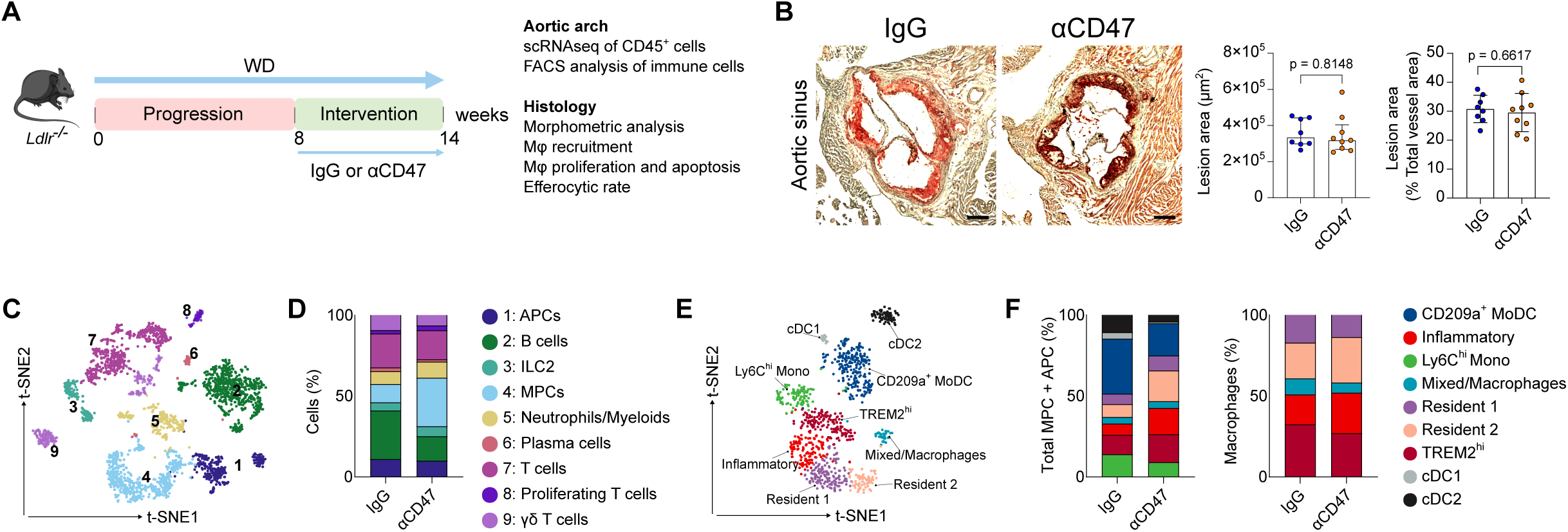
CD47 blockade induces myeloid reprogramming independent of plaque regression. A, Experimental design of the intervention study in an established atherosclerosis model. *Ldlr^-/-^* mice were fed a WD for 14 weeks and treated with IgG or anti-CD47 antibodies during the last 6 weeks. **B,** Quantification of atherosclerotic lesion area in the aortic roots cross-sections stained with Oil Red O (n=8 IgG; n=9 anti-CD47). Scale bar, 250 µm. **C,** t-SNE plot of CD45^+^ cells showing 9 leukocyte clusters in IgG and anti-CD47 groups. **D,** Proportional representation of each major leukocyte cluster within the CD45^+^ population. **E,** t-SNE plot of the myeloid cell compartment after subsetting and reclustering from the total CD45^+^ leukocyte population. **F,** Proportional representation of identified macrophage and dendritic cells populations within the myeloid compartment (left) and macrophage compartment (right). Data and error bars present mean ± SD for parametric and median ± IQR for non-parametric results. Statistical analysis was performed using unpaired Student’s *t*-test (two-tailed) and Mann-Whitney U test (two-tailed). All data and statistical analysis are provided in Source Data.

**Figure 4.**
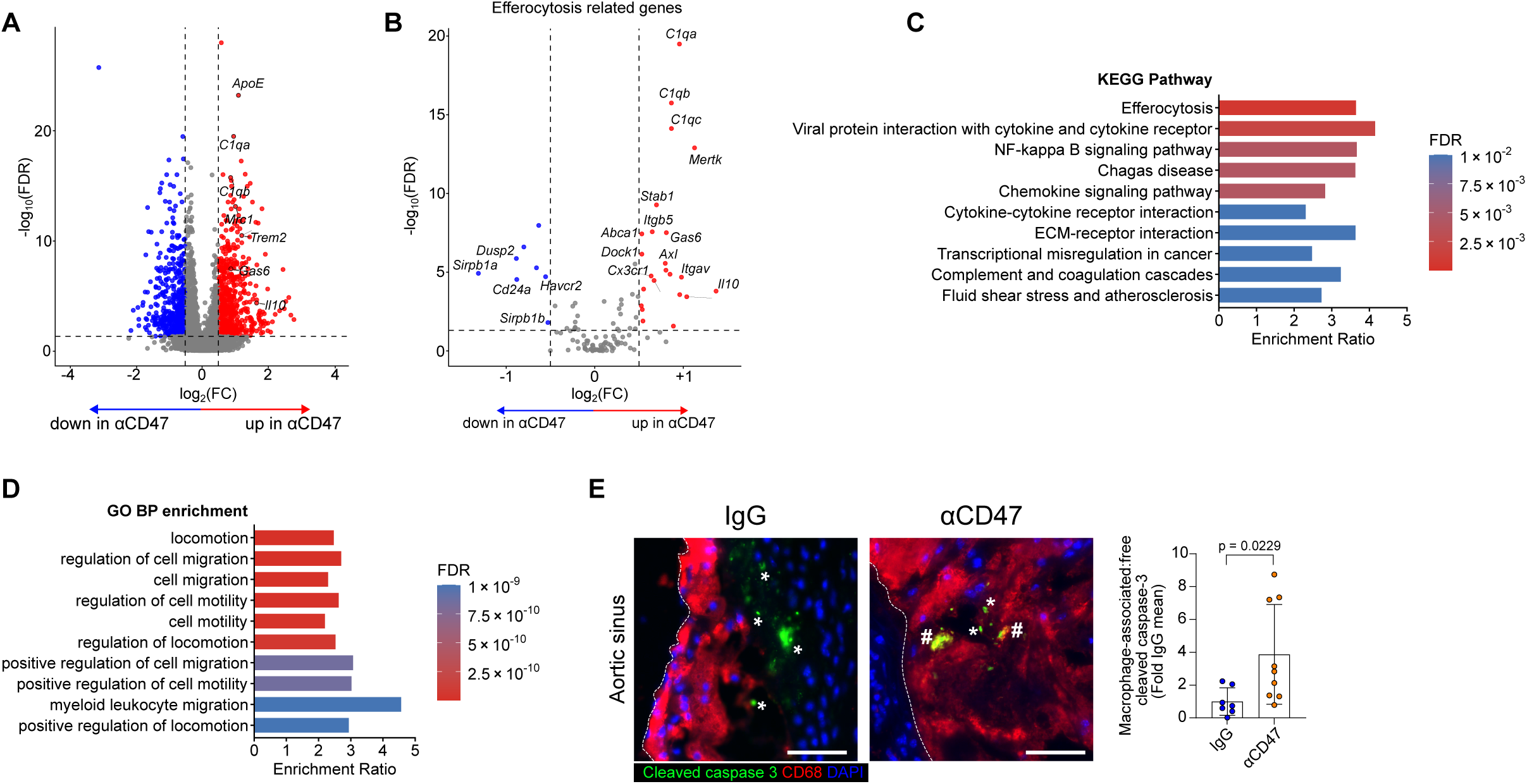
CD47 blockade reactivates efferocytosis machinery in macrophages. **A** and **B,** Volcano plots of differentially expressed genes (DEGs) in myeloid cells following treatment with anti-CD47 versus IgG control. **A,** Global differential expression. **B,** Differential expression of efferocytosis-related genes (KEGG-Pathway, mmu04148). Significant hits (FDR < 0.10; log2FC > 0.5) are color-coded red (upregulated) and blue (downregulated). Canonical efferocytosis genes in (A) are highlighted with black circles. **C** and **D,** Pathway analysis depicting significantly upregulated KEGG pathways (**C**) and biological processes (**D**) in anti-CD47-treated myeloid cells. **E,** Representative immunofluorescence images and quantification of the *in vivo* efferocytosis rate, calculated as the ratio of macrophage-associated (#) to free (*) cleaved caspase-3^+^ apoptotic cells in aortic root sections (n=7 IgG; n=9 anti-CD47). The white dashed line depicts neointima. Scale bar, 10 µm. Data and error bars present mean ± SD. Statistical analysis was performed using unpaired Welch’s *t*-test (two-tailed). All data and statistical analysis are provided in Source Data.

### Anti-CD47 Therapy Regulates Monocyte-Macrophage Dynamics within the Plaque

To elucidate the cellular mechanisms driving the reduction in plaque inflammation, we investigated monocyte kinetics using dual pulse-labelling with EdU and fluorescent microspheres^35,39,43^. While labelling efficiency was comparable between groups, anti-CD47 therapy markedly impaired the recruitment of inflammatory Ly6C^hi^ monocytes into atherosclerotic lesions (Figure 5A; Figure S6A and S6C). In contrast, the entry of patrolling Ly6C^lo^ monocytes remained unaffected (Figure 5B; Figure S6D). This selective suppression of Ly6C^hi^ monocytes was mirrored by a reduction in aortic and circulating levels, but the Ly6C^lo^ monocyte levels were not changed (Figure 5C). However, we observed a paradoxical accumulation of these cells in the spleen, but not in the bone marrow niche (Figure 5D), confirming altered trafficking dynamics rather than systemic depletion. Of note, no differences were observed in other myeloid cell subsets, such as neutrophils, macrophages or dendritic cells, in either the aorta or blood (Figure S6B). To definitively rule out survival defects as a cause for reduced recruitment, we assessed the expression of the M-CSF receptor (CD115), a critical driver of monocyte viability. We observed no reduction in surface CD115 expression on Ly6C^hi^ monocytes, yet the absolute frequency of CD115^+^ Ly6C^hi^ cells increased in peripheral reservoirs (Figure S6E). This accumulation, coupled with intact survival receptor expression, suggests that the therapy limits entry via impaired trafficking rather than inducing cell death. Despite reduced Ly6C^hi^ monocyte recruitment, total macrophage burden (CD68^+^ area) remained unchanged, suggesting that macrophage numbers are maintained through compensatory mechanisms. Notably, local macrophage proliferation (Ki67^+^ CD68^+^) was significantly decreased in anti-CD47-treated animals (Figure 5E and 5F; Figure S6F and S6G)^44^. Supporting a trafficking defect, anti-CD47 treatment downregulated the chemokine and adhesion receptor CX3CR1 specifically on Ly6C^hi^ monocytes in the blood and the spleen, but not in the bone marrow (Figure 5G; Figure S6H). As CX3CR1 mediates endothelial adhesion and crawling, its downregulation, in the absence of changes to survival receptor CD115, suggests that therapy limits monocyte entry by impairing trafficking potential rather than inducing cell death. Furthermore, CellChat analysis of the intervention dataset identified a specific disruption in the *Ccl2-Ccr2* chemotaxis axis within the plaque (Figure 5H–5J; Figure S7A and S7B)^40,45^, driven by the simultaneous downregulation of the ligand in “Resident-2” macrophages and the receptor in TREM2^hi^ macrophages (Figure 5K; Figure S7C and S7D). Additionally, we detected a selective upregulation of the Gas6 pathway within the TREM2^hi^ macrophage cluster, suggesting a reactivation of efferocytosis (Figure S7E). Finally, to evaluate whether the cellular targets of this therapeutic reprogramming are conserved in human disease, we analyzed scRNA-seq data from human coronary atherosclerotic plaques^38^ (Figure S8A and S8B). Cross-species transcriptional mapping revealed that human TREM2^hi^ macrophages, which natively displayed the highest efferocytic activation scores (Figure S8C), were preferentially enriched for the efferocytosis- and survival-associated gene signatures induced by CD47 blockade in our murine models (Figure S8D–S8F). Together, these data demonstrate that CD47 blockade resolves plaque inflammation through two complementary mechanisms: systemically, by impairing Ly6C^hi^ monocyte trafficking and disrupting local CCL-CCR2 recruitment signaling; and locally, by reprogramming existing macrophages toward a pro-efferocytic, reparative state. Transcriptional mapping further reveals that the macrophage program reactivated by this therapy is conserved in human coronary plaques, supporting its translational relevance as a therapeutic mechanism.

**Figure 5.**
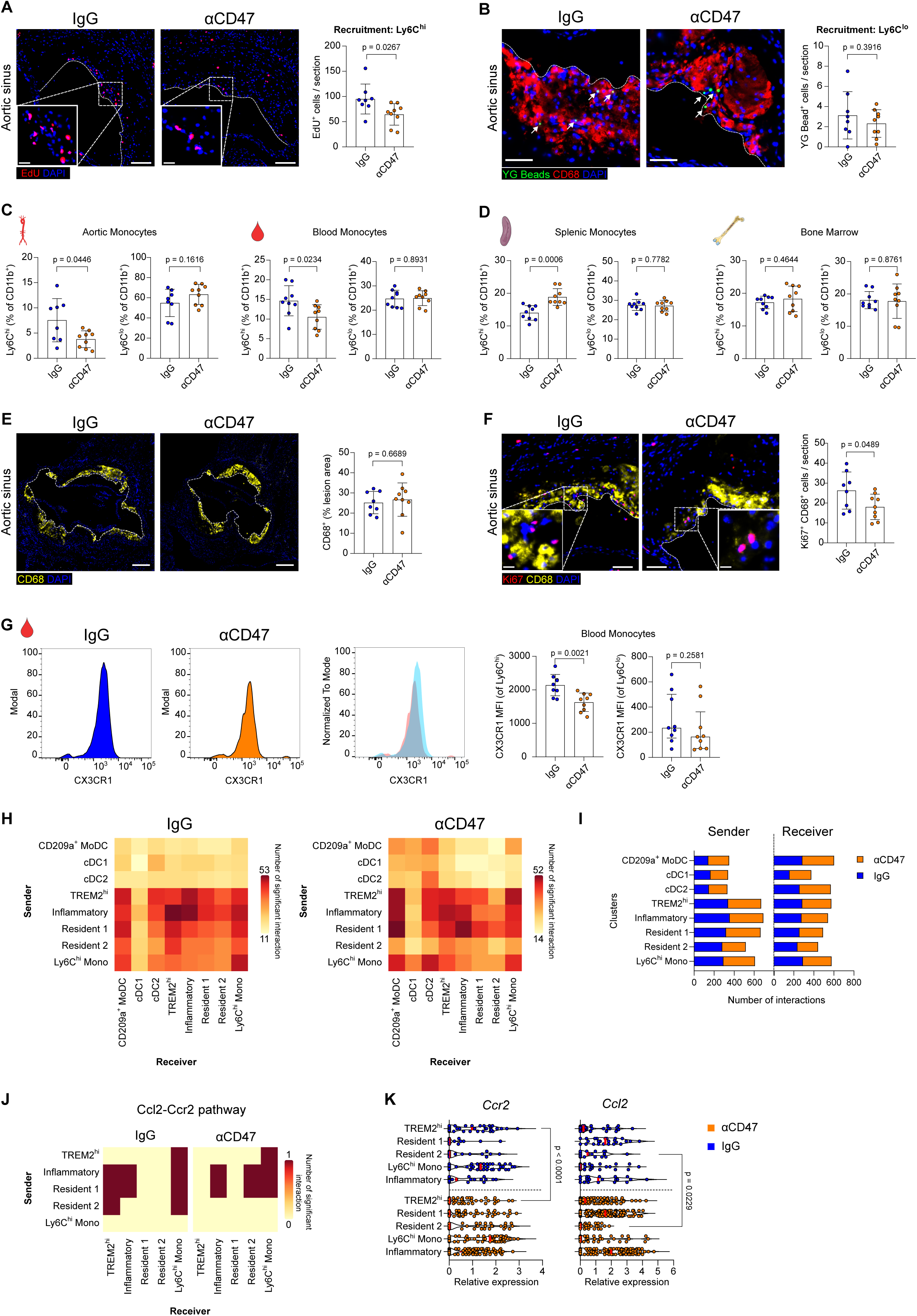
CD47 blockade alters monocyte-macrophage dynamics and innate immune cell communication in atherosclerotic lesions. **A** and **B,** Representative immunofluorescence images and quantification of EdU-labeled Ly6C^hi^ (**A**) and YG fluorescent bead-labelled Ly6C^lo^ (**B**) monocytes (n=8 IgG, n=9 anti-CD47). The white dashed line depicts neointima. Scale bars, 100 µm (**A**) and 50 µm (**B**). **C**, Flow cytometric analysis of Ly6C^hi^ and Ly6C^lo^ monocytes in the aorta (left) and blood (right). **D**, Flow cytometric analysis of Ly6C^hi^ and Ly6C^lo^ monocytes in the spleen (left) and bone marrow (right). **E**, Representative immunofluorescence images and quantification of CD68^+^ macrophages in aortic root sections (n=8 IgG; n=9 anti-CD47). The white dashed line depicts intima and neointima. Scale bar, 250 µm. **F**, Representative immunofluorescence images and quantification of proliferating Ki67^+^ CD68^+^ macrophages in aortic root sections (n=8 IgG; n=9 anti-CD47). The white dashed line depicts neointima. Scale bar, 50 µm. **G**, Quantification of CX3CR1 expression, reported as mean fluorescence intensity (MFI), by flow cytometry on peripheral Ly6C^hi^ and Ly6C^lo^ monocytes (n=8 IgG; n=9 anti-CD47). **H,** CellChat analysis of innate immune cell communication. The heat map depicts the number of interactions per immune cell cluster acting as either sender (ligand) or receiver (receptor). **I**, Number of interactions between innate immune cell clusters serving as sender and receivers. **J**, Heat map showing the number of interactions between macrophage clusters involving *Ccl2-Ccr2* signaling. **K**, scRNA-seq-derived expression of *Ccr2* and *Ccl2* across macrophage subclusters. Data and error bars present mean ± SD for parametric and median ± IQR for non-parametric results. Statistical analysis was performed using unpaired Student’s *t*-test (two-tailed), unpaired Welch’s *t*-test (two-tailed) and Mann-Whitney U test (two-tailed). All data and statistical analysis are provided in Source Data.

## Discussion

The present study demonstrates that pro-efferocytic therapy reshapes atherosclerotic inflammation through mechanisms that extend substantially beyond apoptotic cell clearance and lesion regression. Using a model of established atherosclerosis in which CD47 blockade did not alter lesion size, we show that anti-CD47 therapy remodels the plaque immune microenvironment and promotes inflammation resolution by fundamentally reprogramming the monocyte-macrophage axis. Specifically, CD47 blockade selectively suppresses pro-inflammatory monocyte infiltration and lesional macrophage proliferation while reactivating the efferocytosis machinery within the myeloid compartment, a transcriptional program we find to be conserved in human coronary atherosclerosis. Innate immune checkpoint inhibition targeting the CD47-SIRPα axis is under clinical investigation in hematologic malignancies and solid tumors^46^, and has more recently been evaluated in atherosclerotic cardiovascular disease. Despite preclinical evidence supporting plaque regression, the precise immune mechanisms underlying these therapeutic effects have remained incompletely understood; the present findings begin to address this gap.

Recent single-cell analyses of atherosclerotic plaques have identified a specific subset of foamy macrophages characterized by elevated expression of *Trem2*^4,5,47^. These TREM2^hi^ foamy macrophages exhibit a reduced pro-inflammatory capacity compared to other plaque subsets and are essential for apoptotic cell clearance^5,47^. Indeed, *Trem2^-/-^*macrophages display impaired efferocytosis, leading to necrotic core expansion in hyperlipidemic mice^32^. In this context, we observed that CD47 blockade exerts stage-dependent effects on this population. During early atherogenesis, anti-CD47 therapy limited the formation of foamy macrophages and shifted monocyte trajectories toward beneficial phenotypes. We speculate that more efficient clearance of apoptotic foam cells as they form may limit TREM2^hi^ macrophage accumulation during this stage, though the precise recognition mechanisms remain to be defined. In advanced lesions, anti-CD47 therapy stabilized the TREM2^hi^ macrophage population, assessed by both frequency and transcriptional score, without altering total CD68^+^ plaque area, thereby augmenting the aggregate efferocytic capacity of plaque-associated macrophages. Importantly, cross-species transcriptional mapping revealed that the human TREM2^hi^ macrophage population, conserved in coronary atherosclerosis, displayed the highest enrichment for efferocytosis-associated and macrophage-survival gene signatures transcriptionally primed by CD47 blockade in our murine models. These findings are consistent with previous studies identifying TREM2^hi^ macrophages as lipid-handling, metabolically adapted cells that accumulate in advanced atherosclerotic lesions^4,47^. While these data do not imply that pro-efferocytic therapy directly targets TREM2^hi^ macrophages in humans, they identify this population as a conserved cellular state with the transcriptional machinery to respond to restored efferocytosis signaling. This cross-species concordance underscores the translational relevance of this axis and provides a human cellular correlate for the murine therapeutic response.

Beyond local macrophage reprogramming, the resolution of plaque inflammation relies on controlling net monocyte recruitment^48^, macrophage proliferation^49,50^ and macrophage egress^51^. Using monocyte fate-mapping, we investigated whether CD47 blockade mitigates the pro-inflammatory milieu by altering these kinetics. We observed a selective reduction (approximately 35 %; Figure 5A) in the recruitment of inflammatory Ly6C^hi^ monocytes, but not patrolling Ly6C^lo^ monocytes, to lesions following therapy, suggesting that restricting the influx of pro-inflammatory cells is a critical step in initiating inflammation resolution. Mechanistically, CCR2-dependent recruitment of Ly6C^hi^ monocytes is a primary driver of macrophage replenishment and lesional inflammation^9,10,45^. Consistent with this, computational ligand-receptor analysis predicted a reduction in CCL2–CCR2 signaling between resident and TREM2^hi^ macrophages post-therapy, an inference that warrants orthogonal validation. Furthermore, we observed compartment-specific alterations in circulating and splenic, but not bone marrow, Ly6C^hi^ monocytes, including reduced CX3CR1 surface expression. This pattern suggests that anti-CD47 therapy alters the peripheral phenotype and trafficking potential of inflammatory monocytes at sites of monocyte maturation and reservoir function rather than at the level of monopoiesis. One plausible mechanism is re-education of monocytes within the splenic red pulp, where SIRPα-expressing red pulp macrophages engaged by anti-CD47 therapy may alter the local cytokine or lipid milieu encountered by transitioning monocytes; however, the upstream signals linking efferocytic reactivation to this systemic reprogramming require formal investigation. Additionally, macrophage burden is sustained by a balance between local proliferation and cell death^49^. We found that local macrophage proliferation (Ki67^+^ CD68^+^) in established plaques was reduced by ∼30 % following CD47 blockade. This reduction in both monocyte-derived input and local proliferative renewal did not alter net lesional macrophage density, which may be reconciled by enhanced macrophage survival or residency time downstream of pro-efferocytic reprogramming. Alternatively, efferocytic clearance of apoptotic macrophages may offset reduced proliferative renewal, thereby maintaining lesional macrophage density while reducing the inflammatory quality of the infiltrate. Distinguishing between these mechanisms is an important objective for future work.

Although the innate immune reprogramming and inflammation resolution dynamics described here were observed in a well-established Western Diet model of atherosclerosis^30,32,33^, certain limitations warrant consideration. First, our scRNA-seq experiments were performed on pooled aortic leukocytes, yielding a single sequencing library per experimental group, a design consistent with published aortic single-cell atlases^4,34^ and necessitated by the low immune cell yield from individual aortas. However, this approach precludes formal statistical replication at the sequencing level and limits assessment of inter-individual transcriptional variability. Critically, all major transcriptionally defined observations were independently corroborated by flow cytometry, immunofluorescence, qPCR, and functional efferocytosis assays in fully replicated cohorts; these orthogonal validations provide the primary statistical evidence supporting each claim. Multiplexed single-cell approaches with individual-level sequencing replication will be important in future studies. Second, while the Western Diet-fed *Ldlr*^-/-^ model recapitulates metabolic atherosclerosis and has been extensively characterized as a type 2 diabetes-like state^31^, our study did not perform comprehensive metabolic phenotyping beyond circulating lipid measurements and non-fasting glucose. Formal parameters of systemic insulin resistance, such as fasting insulin levels and HOMA-IR, were not assessed and therefore, insulin resistance was not independently established in our cohorts. Third, our investigation focused primarily on the innate immune compartment; adaptive immune populations were not deeply profiled. Given the emerging role of T and B cells in modulating plaque phenotype and inflammation resolution^35,52,53^, future studies are required to determine how pro-efferocytic therapy influences adaptive immunity and whether these responses contribute to the observed atheroprotection. Fourth, CD47 blockade induces erythrocyte clearance and splenomegaly, as known on-target hematological effects that remain a translational challenge for clinical application^24,54^; these systemic effects did not confound the local aortic immune phenotypes reported here, but their management will be essential for therapeutic translation. Fifth, our single-cell transcriptomic analyses were conducted in male mice. Functional, histological, and flow cytometric experiments included both sexes, but a fully powered sex-stratified transcriptional analysis was beyond the scope of the present study. Given the well-established sexual dimorphism in atherosclerosis and macrophage biology^55^, whether the transcriptional reprogramming described here applies equivalently to female mice remains to be formally determined.

In conclusion, our data demonstrate that pro-efferocytic therapy promotes inflammation resolution in established atherosclerosis not only through reactivation of apoptotic cell clearance, but also through coordinated reprogramming of the monocyte–macrophage axis within a chronic pro-atherogenic metabolic environment. By suppressing Ly6C^hi^ monocyte recruitment, reducing local macrophage proliferation, and enriching efferocytic capacity in plaque-associated phagocytes, CD47 blockade is associated with inflammation resolution while maintaining lesional macrophage stability, effects that may be of particular relevance in patients with established disease and residual vascular inflammation despite lipid-lowering therapy. These findings position reactivation of innate efferocytosis as a mechanistically distinct and translationally relevant strategy for addressing residual cardiovascular risk.

## Acknowledgments

We gratefully acknowledge the data storage service SDS@hd supported by the Ministry of Science, Research and the Arts Baden-Württemberg (MWK) and the German Research Foundation (DFG) through grant INST 35/1503-1 FUGG, the Heidelberg University Core Facilities (Deep Sequencing Core Facility, Flow Cytometry Core Facility), and Servier Medical Art (www.smart.servier.com) for providing components of the Figures. We thank David Ibberson for his expert technical support and all members of the Jarr laboratory for discussions and critical input.

## Author Contributions

M.K., M.I., B.B., A.-L.B., L.B., M.M., and K.-U.J conducted experiments and collected and analyzed data. M.K., N.J.L. and K.-U.J. conceptualized and designed experiments, discussed results and interpreted data. M.K. and K.-U.J. designed figures and wrote the manuscript. M.K. directed the study. K.U.J. supervised the study. All authors commented on and approved the paper.

## Supplemental Figures

**Figure S1. Gating strategy for CD45^+^ aortic leukocytes and systemic characterization of the prevention model.**

**A,** Representative flow cytometry gating strategy to isolate viable aortic CD45^+^ cells, excluding BV510-labelled circulating leukocytes (SYTOX^-^ BV510^-^ PerCP-Cy5.5^+^). **B,** Body weight monitoring during the experimental period of the prevention model (n=12 IgG; n=13 anti-CD47). **C** and **D,** Heart (**C**) and spleen (**D**) weight at euthanasia (n=24 IgG; n=24 anti-CD47). **E,** Quantification of total cholesterol (n=22 IgG; n=21 anti-CD47), high-density lipoprotein (HDL) (n=21 IgG; n=19 anti-CD47), low-density lipoprotein (LDL) (n=21 IgG; n=21 anti-CD47), and non-fasting glucose (n=22 IgG; n=21 anti-CD47) in blood. **F,** t-SNE plot of publicly available scRNA-seq data from aortic CD45^+^ leukocytes of mice fed chow diet (CD) and Western Diet (WD) (Cochain et al., 2018). **G**, t-SNE plot of the fully integrated, batch-corrected dataset showing distinct cell clusters.

Data and error bars present mean ± SD for parametric and median ± IQR for non-parametric results. Statistical analysis was performed using two-way ANOVA, Mann-Whitney U test (two-tailed) and unpaired Student’s *t*-test (two-tailed). All data and statistical analysis are provided in Source Data.

**Figure S2. Monocyte subclusters.**

**A,** Monocle3 pseudotime trajectory of monocyte-macrophage populations superimposed on a t-SNE plot (left) and heat map representing the differentiation progression of the top 20 genes along pseudotime and monocyte-macrophage subclusters (right). **B,** Expression of 5 selected marker genes per monocyte subclusters. The color scale represents log-transformed gene expression. **C**, Dot plot depicting 3 selected marker genes used for monocyte subcluster identification.

**Figure S3. Aortic myeloid cell gating strategy and validation of inflammatory gene expression in the prevention model.**

**A,** Representative flow cytometry gating strategy to analyze major myeloid cells in the aorta. Live, single, CD45^+^ cells were pre-gated to identify CD11b^+^ myeloid cells, Ly6G^+^ neutrophils, Ly6C^+^ monocytes, CD11c^+^ dendritic cells, and F4/80^+^ macrophages. **B**, Flow cytometric analysis of Ly6C^hi^ and Ly6C^lo^ monocytes in aorta-draining lymph nodes (n=10 IgG; n=9 anti-CD47) and bone marrow (n=18 IgG; n=19 anti-CD47). **C,** scRNA-seq gene expression of *Nfkb1 and Nfkbiz* in the MPC cluster. **D,** qPCR analysis of cytokines/chemokine expression (*Tnf, Il5, Il13, Il23, Cx3cl1)* and adhesion molecules (*Vcam1, Icam1)* in aortic arch samples (n=7 IgG; n=5-7 anti-CD47).

Data and error bars present mean ± SD for parametric and median ± IQR for non-parametric results. Statistical analysis was performed using unpaired Student’s *t*-test (two-tailed), unpaired Welch’s *t*-test (two-tailed) and Mann-Whitney U test (two-tailed). All data and statistical analysis are provided in Source Data.

**Figure S4. Intervention model cluster representation.**

**A,** Additional Oil Red O (ORO) staining images depicting lesion size analysis (related to Figure 3B). Scale bar, 250 µm. **B**, Expression of 6 selected marker genes per major cell cluster in the intervention scRNA-seq (left) and dot plot depicting 5 selected marker genes used for identification (right). The color scale represents log-transformed gene expression. **C,** Expression of 8 selected marker genes per MPC + APC subclusters in the intervention model (left) and dot plot depicting 5 selected marker genes used for MPC + APC subcluster identification (right). The color scale represents log-transformed gene expression.

**Figure S5. CD47 blockade reactivates efferocytosis machinery in leukocytes.**

**A,** Volcano plot showing DEGs in CD45^+^ leukocytes following treatment with anti-CD47 versus IgG control. Significant hits (FDR < 0.10; log2FC > 0.5) are color-coded: blue indicates downregulation and red indicates upregulation. Canonical efferocytosis-related genes are highlighted with black circles. **B** and **C,** Pathway analysis depicting significantly upregulated KEGG pathways (**B**) and biological processes (**C**) in anti-CD47-treated CD45^+^ leukocytes. **D,** Gene set enrichment analysis (GSEA) of phagocytosis (GO:0006909) in myeloid cells after anti-CD47 treatment. E, Quantification of total cleaved caspase-3 area in aortic root sections, in relation to total vessel area (TVA, left) and lesion area (right).

**Figure S6. CD47 blockade alters aortic and systemic monocyte dynamics.**

**A,** Flow cytometric analysis of YG bead-labeled (top) and EdU-labeled (bottom) Ly6C^+^ blood monocytes 24 hours after injection (n=6 IgG; n=6 anti-CD47). Population frequencies of EdU^+^ and YG Bead^+^ gates are depicted. **B,** Flow cytometric analysis of CD45^+^ leukocytes, CD11b^+^ myeloid cells, Ly6G^+^ neutrophils, F4/80^+^ macrophages and CD11c^+^ dendritic cells in aorta (top) and blood (bottom) in the intervention model (n=8 IgG; n=9 anti-CD47). **C** and **D,** Additional representative immunofluorescence images of EdU-labeled Ly6C^hi^ (**C**) and YG fluorescent bead-labeled Ly6C^lo^ (**D**) monocytes. The white dashed line depicts neointima. Scale bars, 100 µm; inset, 10 µm (**C**) and 50 µm (**D**). **E**, Flow cytometric analysis of CD115^+^ (M-CSF receptor, CSF1R) Ly6C^+^ monocytes (n=6 IgG; n=6 anti-CD47). Population frequencies of CD115^+^ cells are depicted. **F,** Additional representative immunofluorescence images of CD68^+^ macrophages in aortic root sections. Scale bar, 250 µm. **G**, Additional representative immunofluorescence images of proliferating Ki67^+^CD68^+^ macrophages in aortic root sections. The white dashed line depicts neointima. Scale bar, 50 µm; inset 10 µm. **H**, Quantification of CX3CR1 expression by flow cytometry in splenic (top) and bone marrow (bottom) Ly6C^hi^ and Ly6C^lo^ monocytes (n=9 IgG; n=9 anti-CD47). Data are represented as mean fluorescence intensity (MFI).

Data and error bars present mean ± SD for parametric and median ± IQR for non-parametric results. Statistical analysis was performed using unpaired Student’s *t*-test (two-tailed) and Mann-Whitney U test (two-tailed). All data and statistical analysis are provided in Source Data.

**Figure S7. Ligand-receptor interaction analysis.**

**A,** CellChat analysis of innate immune cell communications. The heat map depicts the max interaction strength per cell communication pair, acting as either sender (ligand) or receiver (receptor). **B,** Heat map showing the interaction strengths between macrophage clusters involving *Ccl2-Ccr2* signaling. **C,** CellChat analysis of the overall *Ccr-Ccl* interaction network between MPC-APC subclusters. **D,** Dot plot depicting ligand-receptor interaction strengths of the specific *Ccr-Ccl* pathway. **E,** Heat maps showing the number of interactions (left) and interaction strengths (right) between macrophage clusters involving *Gas6-Axl/Mertk* signaling.

**Figure S8. Efferocytosis-competent macrophage states are conserved in human atherosclerotic plaques.**

**A,** t-SNE plot of human coronary artery cells; the myeloid cell population is highlighted in the box. **B,** t-SNE representation of human coronary myeloid cells after subsetting and reclustering. **C,** Violin plot of AUCell scores for an apoptotic cell burden gene signature across human coronary myeloid cell clusters. **D,** Analysis workflow showing the curation of DEGs in mouse MPC + APC clusters following CD47 blockade and subsequent human ortholog conversion. Converted genes were separated into functional clusters to identify myeloid responders in human atherosclerotic plaques. **E,** Violin plots depicting AUCell scores for anti-CD47-enhanced efferocytosis/phagocytosis (left) and survival/homeostasis (right) gene sets in human coronary myeloid cells. **F,** AUCell score analysis of negative control gene sets, including metabolic, housekeeping, RNA processing, protein folding, and cytoskeleton-related genes.

